# Lamin B loss in nuclear blebs is rupture dependent while increased DNA damage is rupture independent

**DOI:** 10.1101/2025.02.24.639904

**Authors:** Catherine G. Chu, Nick Lang, Erin Walsh, Mindy D. Zheng, Gianna Manning, Kiruba Shalin, Lyssa M. Cunha, Kate E. Faucon, Nicholas Kam, Sara N. Folan, Arav P. Desai, Emily Naughton, Jaylynn Abreu, Alexis M. Carson, Zachary L Wald, Dasha Khvorova-Wolfson, Leena Phan, Hannah Lee, Mai Pho, Kelsey Prince, Katherine Dorfman, Michael Seifu Bahiru, Andrew D. Stephens

## Abstract

The nucleus houses genetic information and functions separate from the rest of the cell. Loss of nuclear shape results in nuclear ruptures. Nuclear blebs are deformations identified by decreased DNA density, while lamin B levels vary drastically. To determine if decreased lamin B levels are due to nuclear rupture, we used immunofluorescence to measure levels of lamin B and emerin, a nuclear envelope protein that enriches to sites of nuclear rupture. We observed that cell types that exhibit decreased levels of lamin B also show an enrichment of emerin in nuclear blebs. Oppositely, in other cell types, nuclear blebs display maintained levels of lamin B1 and showed no emerin enrichment. To determine how nuclear rupture affects DNA damage, we time lapse imaged nuclear rupture dynamics then fixed the same cells to conduct immunofluorescence of γH2AX and emerin. We find that DNA damage levels are higher in blebbed nuclei independent of nuclear rupture. Thus, we confirm that lamin B1 loss in nuclear blebs is due to nuclear rupture and blebbed nuclei have increased DNA damage that is independent of rupture.

**Summary statement (180-200 characters):** We measured lamin B and DNA damage in blebbed nuclei to determine the effect of nuclear rupture. We find that nuclear rupture causes loss of lamin B in nuclear blebs but that increased DNA damage in blebbed nuclei is independent of rupture.

## Introduction

Since the advent of the microscope, nuclear morphology has been used to both diagnose and prognose human diseases including but not limited to cervical, breast, and prostate cancer. Nuclear blebs are a class of abnormal nuclear morphology associated with disease. In prostate cancer, the frequency of nuclear blebbing correlates with higher Gleason scores, a standard measure of disease severity (Helfand *et al*., 2012). Nuclear blebs arise from perturbations to the two major mechanical components of the nucleus: chromatin or lamins (Hobson and Stephens, 2020; Kalukula et al., 2022; Stephens et al., 2018a). Loss of nuclear strength leads to nuclear shape deformations that cause high curvature and nuclear rupture (Stephens *et al*., 2018b; Xia *et al*., 2018). This resulting loss of nuclear integrity has been reported to cause dysfunction via DNA damage, transcriptional changes, and loss of cell cycle control. Despite these observations, the precise causal relationships between nuclear blebbing, rupture, and their cellular consequences remain to be fully elucidated.

The molecular composition of nuclear blebs relative to nuclear rupture has yet to be determined. Our previous study identified that DNA density loss is the most consistent marker of a nuclear bleb relative to the nuclear body across perturbations and different cell types (Bunner et al., 2024; Pujadas Liwag et al., 2025). In contrast, lamin B levels vary widely between cell types and perturbations, challenging the traditional view that lamin B loss determines a nuclear bleb (Jung-Garcia et al., 2023; Nmezi et al., 2019; Stephens et al., 2018b). The mechanisms driving this heterogeneity in lamin B distribution—where some blebs maintain normal levels while others show dramatic reductions—remain unclear. We hypothesized that lamin B depletion occurs specifically during nuclear rupture, a model supported by our previous findings. Through time lapse imaging, we demonstrated that ruptured nuclear blebs exhibited loss of lamin B signal, whereas non-ruptured blebs maintained lamin B levels comparable to the nuclear body (Bunner et al., 2024). Further validation and investigation of this mechanism’s universality are needed.

Nuclear rupture can be monitored by static population analysis using rupture-specific nuclear envelope proteins and dynamic visualization via NLS-GFP during live cell imaging. Following nuclear envelope rupture, BAF rapidly accumulates at the rupture site and recruits emerin, LEMD2, lamin A/C, and ECRT to repair the nuclear envelope (Halfmann *et al*., 2019; Young *et al*., 2020). Recent studies have highlighted emerin as a reliable marker for detecting recent nuclei rupture events (Young *et al*., 2020). During dynamic live cell imaging, NLS-GFP serves as a real-time indicator of nuclear integrity. NLS-GFP is concentrated in the nucleus and will spill into the cytoplasm upon nuclear rupture, but is re-enriched after nuclear envelope healing, which occurs in about 10 minutes (Pho et al., 2023; Young et al., 2020).

Nuclear rupture has been shown to cause cellular dysfunction most consistently via increased DNA damage measured by γH2AX or 53BP1 foci (Chen et al., 2018; Denais et al., 2016; Irianto et al., 2016; Pho et al., 2023; Raab et al., 2016; Shah et al., 2021; Stephens et al., 2019b; Xia et al., 2018). However, recent studies have demonstrated that nuclear deformation alone, even without rupture, can induce DNA damage (Shah *et al*., 2021). This raises a critical question about the relative contributions of nuclear blebbing and rupture to DNA damage accumulation, necessitating quantitative comparisons between ruptured and non-ruptured nuclear blebs.

Using a diverse array of mammalian cell lines (mouse and human) we investigated the composition of nuclear blebs using DNA, lamin B, and emerin. First, we conducted population immunofluorescence measures to determine bulk behaviors of these three components in the nuclear bleb relative to the nuclear body. Next, we investigated the hypothesis that lamin B is lost from nuclear blebs due to nuclear rupture by measuring the correlation between levels of lamin B and the known nuclear rupture marker, emerin. We then used NLS-GFP cell lines to track nuclear rupture during time lapse imaging into immunofluorescence of those same cells to measure levels of lamin B, emerin, and DNA damage marker γH2AX. Overall, lamin B1 levels decrease upon nuclear rupture in the nuclear bleb while nuclear DNA damage levels are dependent on nuclear blebbing and not rupture.

## Results

### Nuclear blebs consistently exhibit decreased DNA density while lamin B1 and emerin levels fall into two categories

Nuclear blebs are defined by a > 1 µm protrusion of the nucleus. Our previous work established reliable criteria to distinguish these nuclear blebs from micronuclei through systematic scanning and DNA density measurements (Bunner *et al*., 2024). This same work showed that lamin B levels in the bleb varied widely. To determine the source of lamin B bleb variation, we conducted immunofluorescence on numerous cell types and perturbations to track lamin B levels along with known nuclear rupture marker emerin (Halfmann *et al*., 2019; Young *et al*., 2020).

Mouse embryonic fibroblasts (MEFs) provide a cell line that displays nuclear blebs with normal and decreased levels of lamin B relative to the nuclear body (**Figure 1A and C**). MEF cell treatments and genetic modifications provide the ability to compare wild type (WT) relative to chromatin perturbations or lamin perturbations that increase nuclear blebbing. Treatment of MEF cells with histone deacetylase inhibitor VPA or histone methyltransferase inhibitor DZNep causes chromatin decompaction via increased euchromatin or decreased heterochromatin respectively leading to increased nuclear blebbing and rupture (Kalinin et al., 2021; Stephens et al., 2018b; Stephens et al., 2019b). Genetic knockout of lamin B and knockdown of lamin A in MEFs also increased nuclear blebbing and rupture (Berg et al., 2023; Hatch and Hetzer, 2016; Pho et al., 2024; Vahabikashi et al., 2022; Vargas et al., 2012). Immunofluorescence of DNA, lamin B1 (or lamin B2), and emerin reveals that MEF nuclear blebs are heterogeneously decreased for lamin B while emerin levels can be enriched, similar to, or depleted relative to the nuclear body (**Figure 1A**, **C, D**). Again, DNA density via Hoechst staining remained consistently decreased across cells and perturbations (**Figure 1B**), in agreement with our previous work (Bunner *et al*., 2024). Overall, MEF wild type and perturbations show variable lamin B1 and emerin staining in the nuclear bleb.

**Figure 1.**
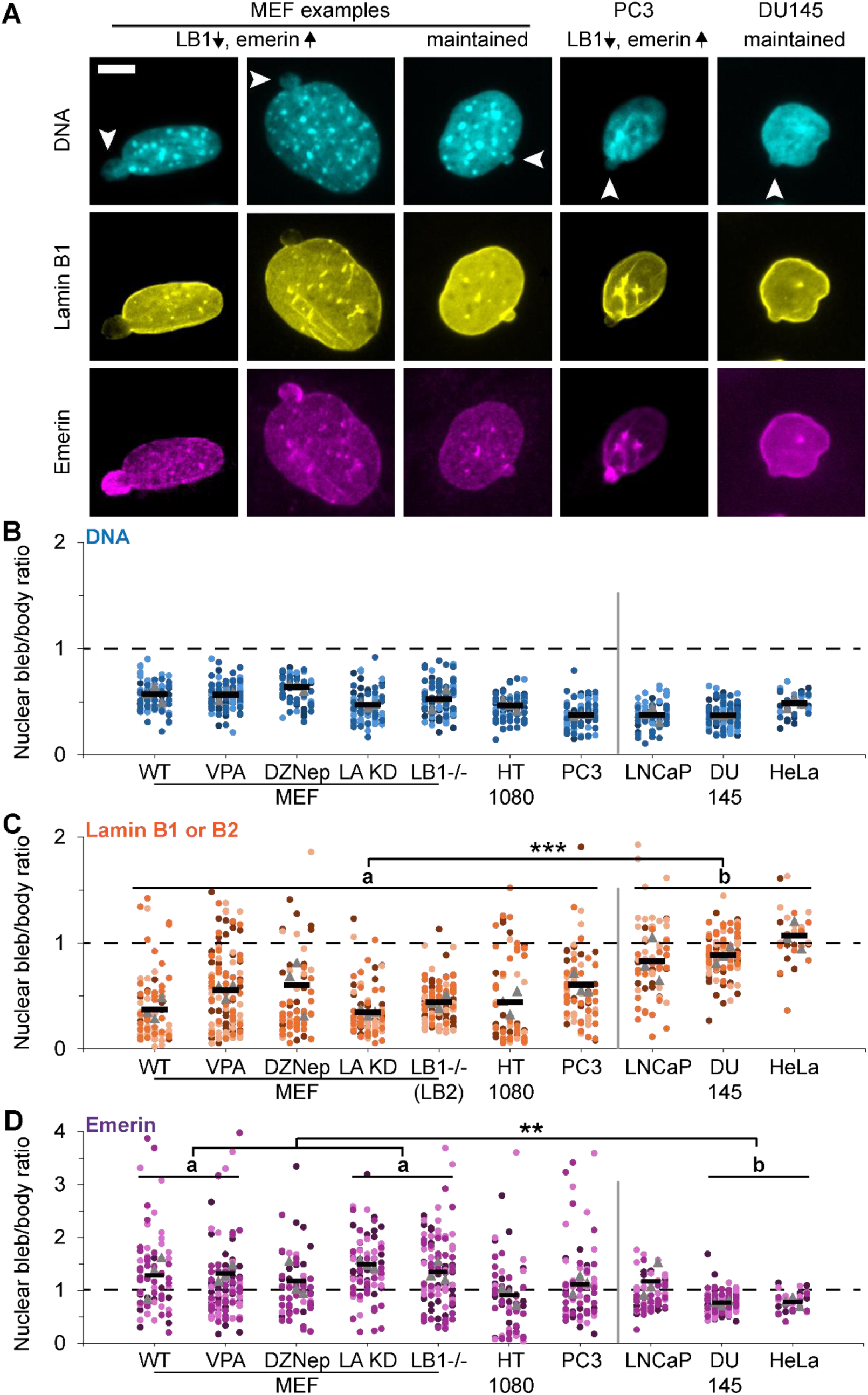
Nuclear bleb vs. body levels reveal consistent loss of DNA while lamin B and emerin levels vary across cell types and perturbations. (A) Example images of a blebbed nucleus in MEF, PC3, and DU145 labeled for DNA via Hoechst (cyan) and immunofluorescence of lamin B1 (yellow) and emerin (magenta). Arrows denote the nuclear bleb. (B-D) Super plots of the relative intensity nuclear bleb/body for (B) DNA, (C) lamin B, and (D) emerin in MEF wild type (WT), chromatin decompaction (VPA and DZNep), and lamin perturbations (LA KD and LB1-/1) and human cell lines HT1080, PC3, LNCaP, Du145, and HeLa. Each condition was done in triplicate with n > 10. Statistical analysis performed using one way ANOVA with Tukey’s post-hoc test with p values reported as * < 0.05, ** < 0.01, *** < 0.001, or no asterisk denotes no significance, p > 0.05. Tables of all comparisons in Supplemental Figure 1. Asterisks in panel C and D similar groups a and b are statistically different for all in the group. N=3-4 where MEF WT, n = 11, 16, 16, 34; VPA n= 38,35,36; DZNep n = 19,10,32; LA KD n = 29, 29 , 27; Lmnb1-/-n = 33, 35, 41; HT1080 n = 18, 15, 29; PC3 n = 16,29,32; LNCaP n = 13,27,27; DU145 n = 37, 40, 35; HeLa n = 11, 10, 10. Mean ± s.e.m is graphed. Scale bar = 10 µm.

Human cancer cell types provide a spectrum between loss or maintenance of lamin B1 in the nuclear bleb relative to the body. Immunofluorescence data reveal that human HT1080 fibrosarcoma and PC3 prostate cancer cell lines showed a MEF-like heterogenous decrease of lamin B1 in the nuclear bleb relative to the nuclear body while emerin levels could be enriched, similar, or depleted in the bleb (**Figure 1C and D**). Oppositely, human cancer cell lines LNCaP, DU145, and HeLa retained similar levels of lamin B1 and emerin in the nuclear bleb compared to the body (**Figure 1C and D**). Thus, depending on the human cell type, nuclear bleb characteristics are significantly different.

Overall, we find two nuclear bleb categories across cell types. MEF, HT1080, and PC3 display a category of nuclear blebs with variably decreased levels of lamin B in the nuclear bleb while LNCaP, DU145, and HeLa display a different category of nuclear blebs that maintain lamin B levels in the nuclear bleb (a vs. b, **Figure 1C**). A portion of these groups also show differential emerin behavior with enriched or maintained levels respectively (a vs. b, **Figure 1D**). This data suggests that lamin B and emerin have an association with nuclear bleb behavior.

### Lamin B and emerin levels are correlated in the nuclear bleb

We hypothesized that nuclear rupture events drive the observed decrease in lamin B1 levels within nuclear blebs. This hypothesis was supported by our previous observation that nuclear blebs maintain lamin B1 levels in the absence of nuclear rupture (Bunner *et al*., 2024). To test this, we measured the correlation of lamin B1 decrease relative to emerin enrichment that occurs at sites of nuclear rupture (Halfmann *et al*., 2019; Young *et al*., 2020). Across MEF wild type and perturbed nucleus conditions nuclear blebs show a heterogenous decrease of lamin B1 relative to the body (**Figure 2A**). Each conditions’ nuclear bleb composition was graphed as lamin B1 vs. emerin bleb/body ratio (**Figure 2C-L**). In 90% of nuclear blebs that maintained lamin B1 levels similar to those in the nuclear body, emerin levels were also similar in the nuclear bleb relative to the nuclear body (**Figure 2C-F**). In contrast, blebs with decreased lamin B1 frequently showed substantial emerin enrichment, often several-fold higher than the nuclear body. However, not all lamin B1 decreased nuclear blebs showed emerin enrichment as some measured similar or reduced emerin in the nuclear bleb relative to the body. Overall, we find that nuclear rupture marker emerin does not enrich in nuclear blebs that maintain lamin B1, while blebs with decreased lamin B1 are frequently enriched in emerin.

**Figure 2.**
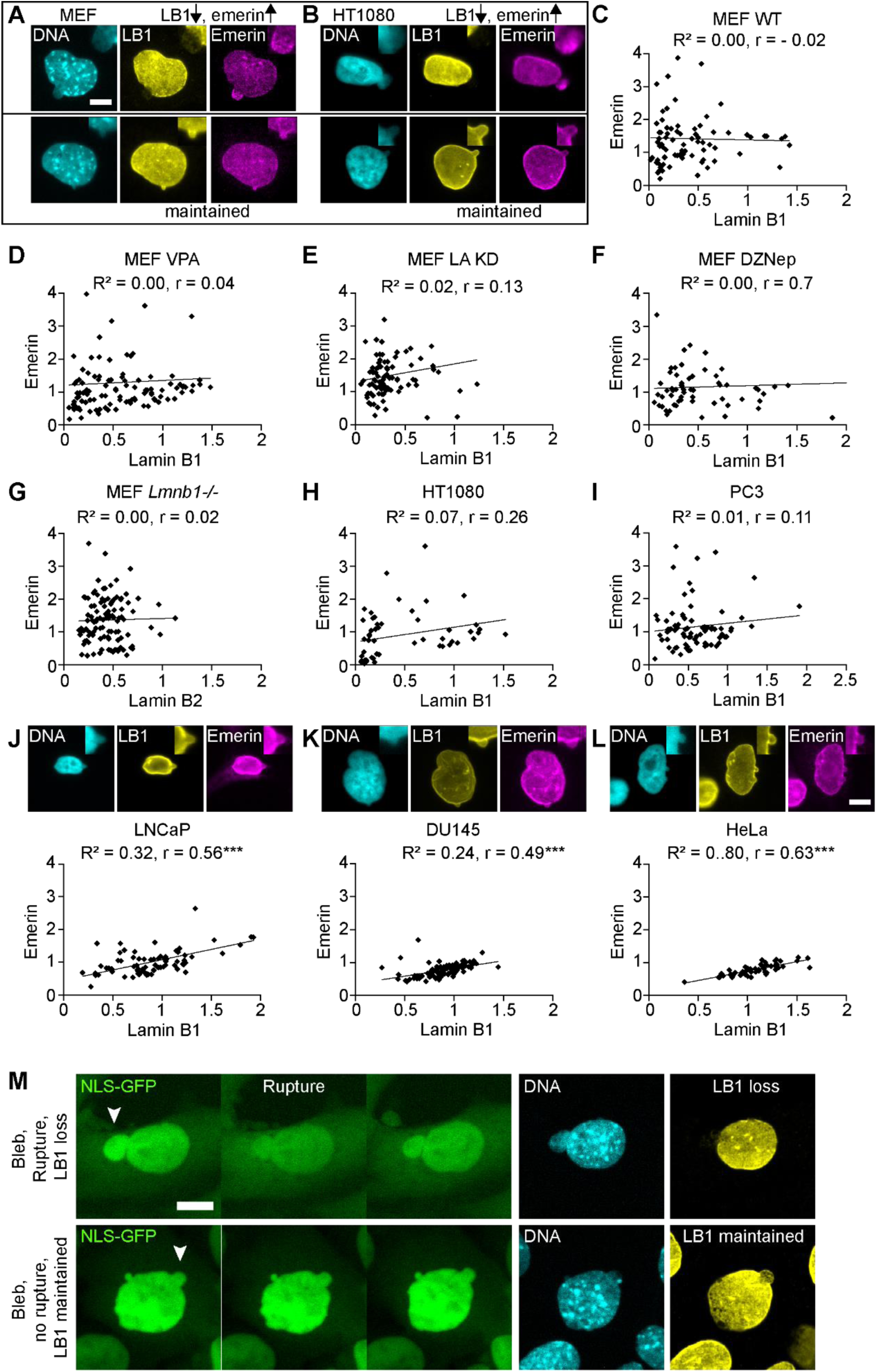
Nuclear blebs with decreased lamin B1 display increased emerin while maintained lamin B1 shows correlation with emerin levels. (A-B) Example images of nuclear blebs with or without changes in lamin B1 and emerin for (A) MEF and (B) human HT1080 cell lines. Top right insets show a closeup of the bleb. (C-L) Quantitative analysis of nuclear bleb/body ratios comparing lamin B1 versus emerin levels across cell lines: MEF wild type (WT, n=79), MEF VPA (n=110), MEF LA KD (n=85), MEF DZNep (n=61), MEF Lmnb1-/- (n=109), HT1080 (n=53), PC3 (n=77), LNCaP (n=79), Du145 (n=112), and HeLa (n=31). Cell lines (A) MEF and (B) HT1080 show nuclear blebs with decreased lamin B1 and increased emerin while cell lines (J) LNCaP, (K) Du145, and (L) HeLa that maintain lamin B1 levels in the bleb show a linear relation of emerin nuclear bleb to body ratio around 1. Each set of data was fit for linear regression denoted by R^2^ value and statistically significant correlation via Pearson’s correlation (*r*). (J, K, L) Example images along with graphs of emerin relative to lamin B1 of LNCaP, Du145, and HeLa nuclear blebs stained for DNA via Hoechst (cyan) and immunofluorescence of lamin B1 (yellow) and emerin (magenta) that largely maintain similar levels in the nuclear bleb and body and have a strong Pearson’s correlation (*r*). Top right insets show a closeup of the bleb. (M) Example images of a time lapse NLS-GFP imaging into immunofluorescence via Hoechst DNA stain (cyan) and lamin B1 (yellow) of a nuclear bleb that does and does not rupture. Scale bar = 10 µm.

We next investigated human cell lines presenting heterogeneously decreased lamin B levels in the nuclear bleb to determine if this phenomenon is reproducible in other cell types. Human cancer cell line HT1080 is well reported to have nuclear blebs that are rupture prone (Stephens et al., 2019b). Upon analysis of HT1080 nuclear blebs, we found a relationship between lamin B1 and emerin similar to that observed in MEFs (**Figure 2B**). For blebs that have a similar level of lamin B1 in the bleb and body (bleb/body ratio =1), emerin levels in the nuclear bleb and body were also similar (**Figure 2H**). Decreased lamin B1 in nuclear blebs of HT1080 nuclei were associated with both emerin enrichment and reduction. This trend was also present in prostate cancer cell line PC3 (**Figure 2I**) and previously reported in human fibroblast cells (Janssen et al., 2022). These findings recapitulate that bleb-based rupture prone nuclei display loss of lamin B1 in conjunction with enrichment of emerin due to nuclear rupture.

Human cell lines LNCaP, DU145, and HeLa provide the opportunity to investigate nuclear blebs that retain lamin B1 relative to the body (**Figure 1 and Figure 2J-L**). Correlation graphs of lamin B1 and emerin levels in LNCaP, DU145, and HeLa show rare enrichment of emerin suggesting these nuclear blebs rupture infrequently if at all. Interestingly, the relationship between lamin B1 and emerin levels in the bleb follows a linear correlation, suggesting a coupled and slight variation of the nuclear lamina/envelope present. These levels are also centered around 1 or similar levels relative to the rest of the nucleus. Thus, nuclear blebs that maintain normal lamin B1 levels consistently show normal emerin distribution without enrichment, indicating an absence of nuclear rupture events.

To verify that nuclear lamin B levels reflect nuclear rupture we did time lapse imaging into immunofluorescence of lamin B levels in MEF cells. Nuclear blebs that rupture, determined by NLS-GFP spilling, showed a drastic loss of lamin B in the nuclear bleb relative to the body (**Figure 2M**, top). Oppositely, nuclear blebs that did not rupture throughout the 3 hours of time lapse imaging maintained lamin B immunofluorescence signal in the bleb (**Figure 2M**, bottom). These data agree that when lamin B1 is maintained emerin levels are not enriched because there was no rupture, which agrees with past studies (Bunner et al., 2024)

### Time lapse imaging into immunofluorescence reveals that increased DNA damage in nuclear blebs is rupture independent

We imaged cells at two-minute intervals for 3 hours and then immediately conducted immunofluorescence to connect nuclear blebbing and rupture dynamics with protein content of the nucleus and bleb (**Figure 3A**). Previous studies of immunofluorescence report that blebbed nuclei have higher levels of DNA damage than normally shaped nuclei (Pho et al., 2023; Stephens et al., 2019b). However, the mechanism for increased DNA damage remains under debate. Numerous publications, with or without data, hypothesize that nuclear ruptures cause increased DNA damage (Kalukula et al., 2022; Stephens, 2020). To test this hypothesis, we compared DNA damage levels across normally shaped nuclei (non-blebbed and no rupture), blebbed nuclei that rupture, and blebbed nuclei that did not rupture during the time lapse. Normally shaped nuclei maintained a low level of DNA damage relative to blebbed nuclei (**Figure 3A-C**), recapitulating our previous static studies (Pho et al., 2023; Stephens et al., 2019b). Unexpectedly, blebbed nuclei showed similarly elevated DNA damage levels regardless of rupture status (**Figure 3B and C**). This finding reveals that nuclear blebbing is sufficient to induce increased DNA damage at levels comparable to those observed in ruptured nuclei.

**Figure 3.**
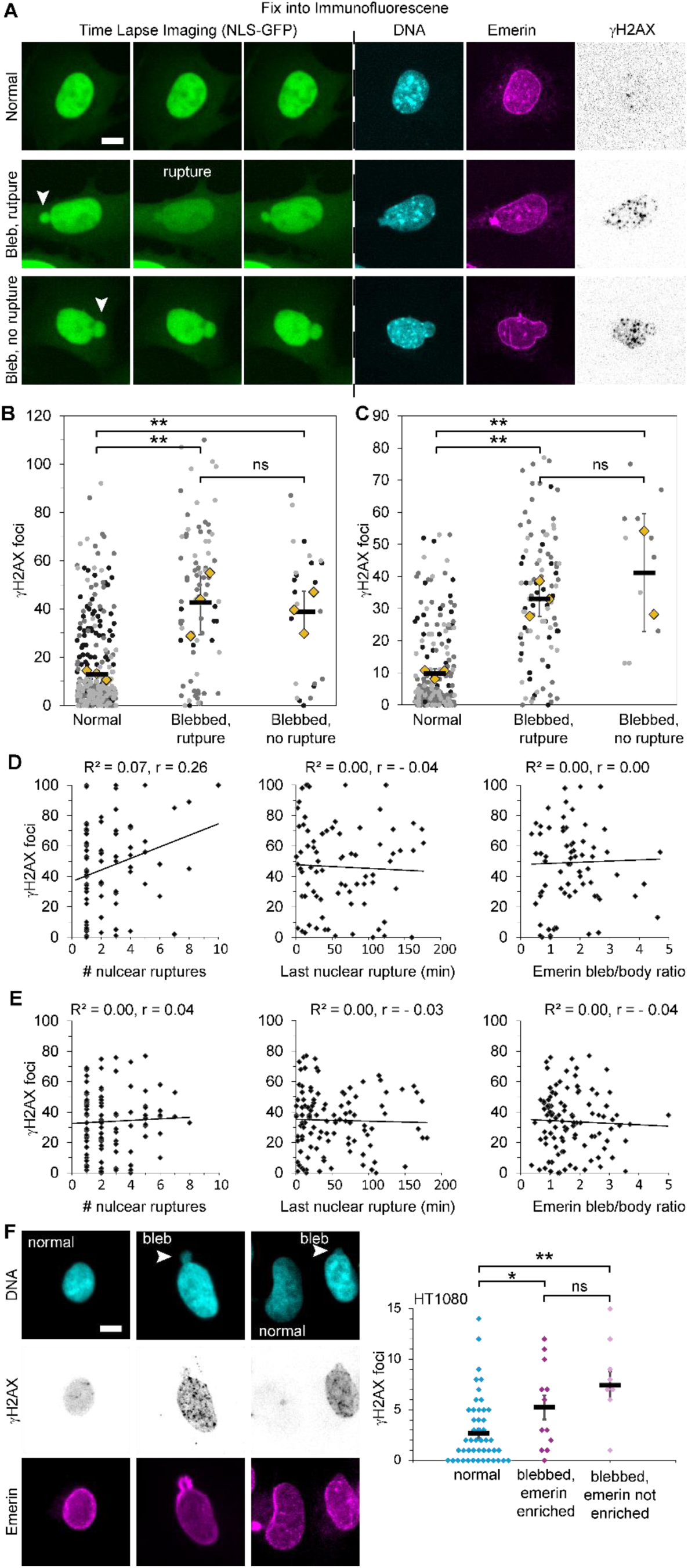
DNA damage via γH2AX increases in blebbed nuclei independent of nuclear rupture. (A) Example images of MEFs from time lapse imaging via NLS-GFP and following immunofluorescence of the same nucleus stained for DNA via Hoechst (cyan), emerin (magenta), and DNA damage foci labeled by γH2AX. (B-C) Super plots of the number of DNA damage foci measured via γH2AX for (B) MEF WT or (C) MEF VPA-treated in nuclei that are normally shaped, blebbed and rupture, and blebbed and no rupture (MEF WT normal n = 186, 173, 112; bleb rupture n = 9, 44, 26; bleb no rupture n = 11, 5, 8; MEF VPA normal n = 53, 120, 56; bleb rupture n = 29, 42, 32; bleb no rupture n = 7, 5). (D-E) Scatter plot of (D) MEF WT or (E) MEF VPA-treated in nuclei γH2AX foci per number of nuclear ruptures, time since last nuclear rupture, and emerin bleb/body ratio along with R2 and r Pearson correlation. (F) Example images of HT1080 nuclei normally shaped or blebbed stained for DNA via Hoechst (cyan), emerin (magenta), and DNA damage foci labeled by γH2AX (n = 47, 24, 7 nuclei respectively). Two-tail unpaired Student’s t-test p values reported as * < 0.05, ** < 0.01, *** < 0.001, no asterisk denotes no significance, p > 0.05. Mean ± s.e.m is graphed. Scale bar = 10 µm.

Given our finding that nuclear blebs, rather than rupture events, drive DNA damage accumulation, we investigated how other rupture-associated metrics correlate with DNA damage levels. In both wild-type MEFs and VPA-treated cells (chromatin decompaction), we found no correlation between DNA damage levels and number of nuclear ruptures (**Figure 3D and E**). Similarly, neither the time elapsed since the last nuclear rupture nor the degree of emerin enrichment showed a correlation with DNA damage levels. (**Figure 3D and E**). Taken together, the frequency and recency of nuclear rupture do not significantly impact increased levels of DNA damage associated with nuclear blebs.

To determine the generality of this phenomenon, we used human cancer cell line HT1080. Consistent with our previous work, HT1080 cells showed elevated DNA damage foci in blebbed nuclei compared to normally shaped nuclei (Pho et al., 2023; Stephens et al., 2019b). Importantly, DNA damage levels, while elevated in blebbed nuclei, showed no correlation with emerin levels in nuclear blebs, a marker of recent nuclear rupture (**Figure 3F**). Taken together across cell types, increased levels of DNA damage are associated with blebbed nuclei independent of nuclear rupture.

### LNCaP blebbed nuclei lack rupture but have increased DNA damage

LNCaP nuclei provide the possibility to explore DNA damage levels in nuclear blebs that might not rupture. LNCaP nuclei have maintained lamin B levels in their nuclear blebs and show little to no emerin enrichment (**Figure 2J**). We imaged LNCaP cells transiently transfected with NLS-GFP and found that no imaged nuclear blebs ruptured over 22.5 hours of imaging **(Figure 4A**, n = 11 blebbed nuclei). Next, we assayed normal and blebbed nuclei for increased DNA damage in nuclear blebs. We found that blebbed nuclei, in which we found no nuclear ruptures, had increased levels of DNA damage relative to normally shaped nuclei (**Figure 4**, **B and C**). In a cell type that presents no nuclear ruptures and maintained levels of lamin B1, blebbed nuclei still had higher levels of DNA damage compared to normally shaped nuclei. Thus, data from both nuclei that rupture and nuclei that do not rupture supports that increased DNA damage is due to nuclear blebbing and not due to nuclear rupture.

**Figure 4.**
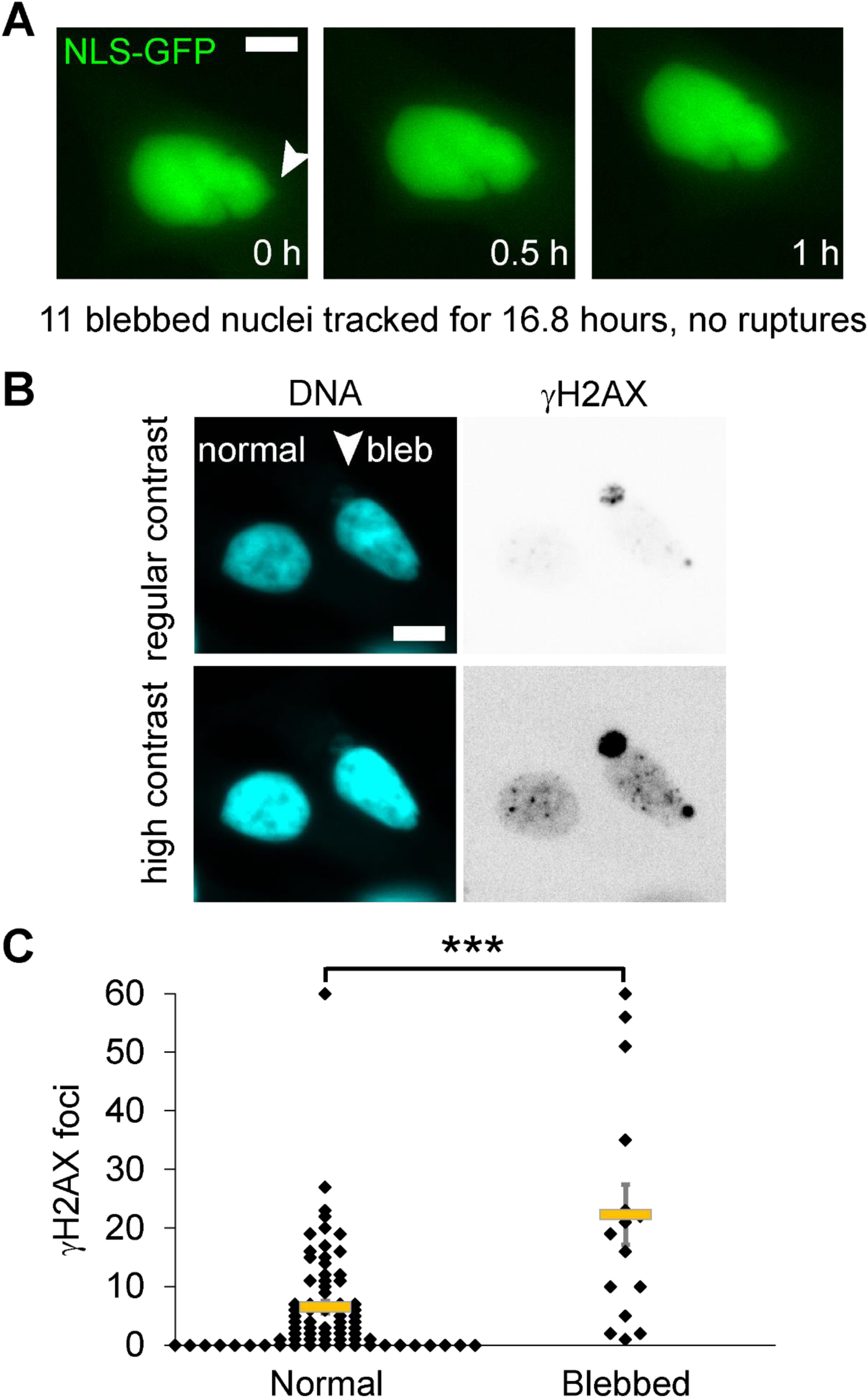
LNCaP blebbed nuclei do not show nuclear rupture but have increased DNA damage. (A) Example images of LNCaP blebbed nucleus that does not show loss of NLS-GFP during live cell imaging (two biological replicates, total 11 blebbed nuclei imaged for 22.25 hours showed 0 nuclear rupture). (B) Example images of DNA labeled via Hoechst (cyan) and γH2AX DNA damage maker foci (inverted gray scale) shown in regular and high contrast to help visualize the bleb. (C) Graph of normal elliptical shaped and blebbed LNCaP nuclei DNA damage foci marked by γH2AX (normal n = 75 and blebbed n = 15). Two-tail unpaired Student’s t-test p values reported as * < 0.05, ** < 0.01, *** < 0.001, no asterisk denotes no significance, p > 0.05. Mean ± s.e.m is graphed. Scale bar = 10 µm.

## Discussion

Nuclear bleb lamin B composition is an indirect marker of bleb integrity, challenging previous studies that incorrectly defined nuclear blebs based solely on lamin B levels. We show that DNA density levels are the most consistent marker of a nuclear bleb (**Figure 1B**) in strong agreement with our previously published work (Bunner et al., 2024; Pujadas Liwag et al., 2025). Furthermore, we clarify that lamin B1 presence or absence is determined by nuclear rupture history as evidenced by measurements with nuclear rupture marker emerin and time lapse imaging into immunofluorescence (**Figures 2 and 3**). Nuclear blebs are significant features of cellular pathology, contributing to nuclear dysfunction that may promote disease progression (Kalukula et al., 2022; Stephens et al., 2019a). We provide novel data showing that the presence of a nuclear bleb independent of nuclear rupture denotes dysfunction (**Figures 3 and 4**). This is a paradigm shift from the assertion that nuclear rupture was the main cause of dysfunction. These results both verify why lamin B levels in the nuclear bleb are variable between cell types and reveal that the presence of a nuclear bleb alone is a sufficient marker of underlying nuclear dysfunction.

### Lamin B1 loss in nuclear blebs indicates prior rupture

Previous studies incorrectly suggested that lamin B1 loss defines a nuclear bleb. Our data clarifies instead that loss of lamin B1 is an indirect marker for past nuclear bleb rupture. Lamin B1 loss in the bleb is cell type and rupture dependent measured by both direct visualization of NLS-GFP spilling and correlation with rupture marker emerin (**Figure 2**). Other studies have detailed the correlation between loss of lamin B1 and emerin enrichment in the nucleus which is reliant on BAF recruitment of emerin (Janssen et al., 2022). Furthermore, lamin B1 loss does not affect nuclear blebbing frequency compared to either chromatin or lamin perturbations (Berg et al., 2023; Pho et al., 2023; Stephens et al., 2018b). Our findings are also consistent with laser ablation studies where lamin B is lost upon ablation of the nuclear envelope while lamin A and lamin C remain (Kono et al., 2022). This selective loss likely results from lamin B’s unique integration into the nuclear envelope via its farnesylation group, which is absent in lamins A and C (Butin-Israeli et al., 2012). The loss of envelope-associated proteins following rupture aligns with previous observations (Hatch and Hetzer, 2016). These results resolve a key misconception in the field, by revealing that lamin B1 levels are representative of nuclear bleb integrity history.

Notably, we observed a subset of nuclear blebs showing concurrent reduction in both lamin B1 and emerin levels. Interestingly, a subpopulation of nuclear blebs with reduced lamin B1 levels also showed reduced emerin. This finding suggests potential permanent damage to the nuclear lamina in these blebs. We hypothesize that while these blebs may have restored envelope compartmentalization, they failed to fully reconstruct their lamina structure. Further investigation is needed to determine the specific consequences of this disrupted lamina architecture.

### Nuclear blebs signal dysfunction independent of rupture

Previous studies proposed that compromised nuclear mechanics leads to deformation and subsequent rupture as the primary pathway to cellular dysfunction. Specifically, loss of nuclear integrity is thought to cause dysfunction through diffusion of nuclear proteins out of the nucleus and cytosolic proteins into the nucleus (Denais et al., 2016; Raab et al., 2016; Xia et al., 2018). DNA damage emerged as the most frequently reported dysfunction associated with altered nuclear mechanics and morphology. Our previous work demonstrated both increased DNA damage in blebbed nuclei and frequent rupture of nuclear blebs (Pho et al., 2023; Stephens et al., 2019b). However, our new data reveal that nuclei with nuclear blebs that did not rupture have a similar increase in DNA damage to nuclei that have ruptured (**Figures 3 and 4**). This finding causes a paradigm shift in the dominant mechanism of DNA damage.

Nuclear dysfunction in a blebbed but unruptured nucleus could stem from mechanical and/or structural features. Supporting evidence for deformation-driven DNA damage exists in earlier literature. In one of the first papers to report that nuclear rupture causes DNA damage, the example image shows that deformation results in increased DNA damage foci well before the nucleus ruptures (Denais et al., 2016). Increased DNA damage has also been reported in nuclei that are mechanically deformed but not ruptured, reportedly due to DNA replication stress (Shah et al., 2021). One proposed mechanism could be that nuclear deformation is separating repair factors from their needed site of function (Irianto et al., 2016). This finding is further supported by work showing that glioblastoma cell types U251MG and U87MG both have nuclear blebs but the more replication active U251 shows more DNA damage (Kamikawa et al., 2023). This observation aligns with other reports showing that DNA damage in blebbed and ruptured nuclei is due to DNA replication stress (Shah et al., 2021; Xia et al., 2019). These findings collectively suggest that the combination of active DNA replication and mechanical stress in blebbed nuclei, rather than rupture alone, drives DNA damage accumulation.

### Nuclear blebs as a site of genome instability

The nuclear bleb might be a site of increased genome instability that drives a more aggressive disease state. Nuclear blebs are diagnostic markers of disease progression in prostate cancer (Helfand *et al*., 2012). Our studies of LNCaP cells, an early-stage prostate cancer model, reveal nuclear blebs that maintain lamin B1 and emerin levels without rupture (**Figures 1,2**, **and 4**). However, blebbed LNCaP nuclei show increased DNA damage with markers in both the nuclear body and the bleb. Progression of prostate cancer has also been linked to DNA translocations in androgen responsive genes. LNCaP cells treated with DHT increase nuclear blebbing and activation of androgen responsive genes that disproportionately end up in the nuclear bleb (Helfand *et al*., 2012). Our other published work shows the importance of transcription activity in causing nuclear blebbing (Berg et al., 2023). Thus, increased transcriptional activity aids nuclear blebbing which causes transcriptionally active chromatin to move into the nuclear bleb, a separate environment that likely exacerbates genome instability, independent of nuclear rupture.

LNCaP, PC3, and DU145 are progressively aggressive models of prostate cancer that all exhibit nuclear blebs. LNCaP, an early model of prostate cancer, shows maintenance of lamin B1 and emerin similar to the most aggressive model DU145 while intermediate model PC3 shows loss of lamin B1 and emerin enrichment. The major changes between LNCaP and PC3 remain unclear, which suggests that rupture is not essential to disease progression. Alternatively, nuclear rupture is not required but does occur later in disease progression. In this scenario, unruptured LNCaP nuclei still incur dysfunction due to their nuclear blebs. Upon becoming more aggressive like PC3, nuclear rupture occurs further driving disease progression. Finally, DU145 cells, with their distinctive nuclear morphology, may represent a state where rupture becomes dispensable or where disease progression follows alternative mechanisms.

Paradoxically, the absence of rupture might exacerbate cellular dysfunction. Specifically, the lack of rupture would stop the enrichment of BAF, emerin, LEM2, and ESCRT. BAF and emerin have important chromatin interaction roles that upon their local enrichment in a ruptured bleb, aid compaction and reorganization (Alfert et al., 2019; Marano and Holaska, 2022) while general loss of emerin can lead to nuclear invasiveness (Hansen et al., 2024). In conclusion, the absence of rupture could remove possible points in which the nuclear environment could be repaired through multiple different mechanisms.

Future work should aim to determine the underlying cause of dysfunction in both nuclei with blebs and the microenvironment of the nuclear bleb. Does the mechanical state of having your nucleus herniate to form a bleb suggest the genome is under physical stress to cause dysfunction? While one study found that compression with AFM can cause increased DNA damage, more and differing experiments will are needed to assess this possibility. For example, micropipette aspiration (Irianto et al., 2016; Zhang et al., 2019), nuclear confinement (Le Berre et al., 2012; Nader et al., 2021), experiments with single micropipette micromanipulation (Neelam et al., 2015) or dual micropipette micromanipulation (Currey et al., 2022) could apply different types and modify global vs. local deformations. Does the nuclear bleb have a different or separate microenvironment? While we have reported that the nuclear bleb has consistently decreased DNA density (Bunner et al., 2024) and destabilized packing domains (Pujadas Liwag et al., 2025), future studies of chromatin dynamics, protein diffusion, and in situ chromosome track would greatly aid determining the nuclear bleb environment.

## Materials and Methods

### Cell culture

Mouse Embryonic Fibroblast (MEF) were previously described in (Shimi *et al*., 2008; Stephens *et al*., 2018b; Vahabikashi *et al*., 2022). MEF wild-type (WT), lamin A knockdown (LA KD), and lamin B1 knockout (*Lmnb1-/-*) cells were cultured in DMEM (Corning) completed with10% fetal bovine serum (FBS; HyClone) and 1% Penicillin/streptomycin (PS; Corning), incubated at 37°C and 5% CO2. Cells were passaged after reaching 80-90% confluency or every 2 to 3 days. Cells were trypsinized, replated, and diluted with DMEM (Corning). Human fibrosarcoma cell HT1080 cells and HeLa human cervical cancer cells were cultured and passaged similarly. HT1080 and HeLa cells were obtained from ATCC.

Three prostate cancer cell lines were used: LNCaP, DU145, and PC3. DU145 and LNCaP cells were cultured in RPMI 1640 with 10% fetal bovine serum (FBS) and 1% penicillin/ streptomycin. PC3 cells were cultured in DMEM completed with 10% fetal bovine serum (FBS) and 1% penicillin.

### Drug treatments

MEF WT cells were treated with either 4 mM valproic acid (VPA, 1069-66-5, Sigma, (Gurvich et al., 2004)) or 1µM 3-deazaneplanocin (DZNep, Cal Biochem (Miranda *et al*., 2009)) for 24 hours before fixation for immunofluorescence or time lapse imaging. HT1080 and Hela cells were treated with VPA similarly to MEF cell lines.

### Immunofluorescence

Cells were grown on coverslips in preparation. Cells were fixed with 3.2% paraformaldehyde and 0.1% glutaraldehyde in phosphate buffered saline (PBS) for 10 minutes. Between steps cells were washed with PBS Tween 20 (0.1%) and Azide (0.2 g/L) three times, denoted PBSTw-Az. Next, cells were permeabilized by 0.5% Triton-X 100 in PBS for 10 minutes. Again, cells were washed with PBS-Tw-Az. Humidity chambers were prepared using petri dishes, filter paper, sterile distilled water, and parafilm. 50µL of primary antibody solution was applied to the parafilm and the cell-side of the coverslip was placed on top to incubate for 1 hour at 37°°C in the humidity chamber. Primary antibodies used were rabbit anti-Lamin B1 (1:1000, ab16048 Abcam) and mouse anti-emerin (1:2000, NCL-emerin, Leica Biosystems). The humid chambers were removed from the incubator and the coverslips were washed in PBS-Tw-Az. New humidity chambers were made for the secondary incubation. 50µL of secondary antibody was placed on the parafilm, and the coverslips were placed cell-side down. The secondary antibody solution contained goat anti-rabbit 488 (1:200, 4412s, Cell Signaling Technology) and goat anti-mouse 555 (1:200, 4409s, Cell Signaling Technology) and incubated in humidity chambers at 37°C for 30 minutes. After, coverslips were washed with PBS-Tw-Az and placed cell-side down on a slide with a drop of mounting media containing DAPI. Alternatively, cells were stained with a 1 µg/mL (1:10,000) dilution of Hoechst 33342 (Life Technologies) in PBS for 5 minutes, washed with PBS 3 times, and mounted with Prolong Fade gold. Slides were allowed to cure for four days at 4°C before imaging.

For live cell imaging into immunofluorescence experiments, we used gridded imaging dishes (Cellvis #D35-14-1.5GI). Similarly, cells were fixed in the gridded imaging dish with 4% paraformaldehyde for 15 minutes. Next, cells were washed in PBS, permeabilized by 0.1% Triton-X 100 in PBS for 15 minutes, and neutralized by 0.06% Tween 20 for 5 minutes. Again, cells were washed with PBS. Cells were then blocked with 2% Bovine Serum Albumin (BSA) for 1 hour. 500µL of primary antibody solution was applied to the cells at room temperature for 2 hours. Primary antibodies used were either rabbit anti-lamin B1 (1:1000, ab16048, Abcam) or mouse anti-emerin (1:1000, NCL-emerin-A, Leica Biosystems). The cells were again washed with PBS. 500µL of secondary antibody was then applied to the cells at room temperature for 1 hour. Secondary antibodies used were either Alexa Fluor 555 anti-mouse IgG (1:1000, 4410s, Cell Signaling Technology) or Alexa Fluor 647 anti-mouse IgG (1:1000, 4413s, Cell Signaling Technology). After the cells were washed with PBS, 500 µL of conjugate antibody was applied to the cells at room temperature for 2 hours. The conjugate antibody used was γH2AX-647 rabbit (1:1000, 9720, Cell Signaling Technology). Cells were then washed with PBS and stained with 1 µg/mL (1:10,000) dilution of Hoechst (H3570, Invitrogen) in PBS for 15 minutes. Cells were washed and stored in PBS until imaging. Transmitted light images were acquired to align the gridded dish after fixation so that the same cells from the time lapse imaging could be imaged for their respective immunofluorescence of DNA via Hoechst 33342, emerin, Lamin B1, and DNA damage via yH2AX.

### Immunofluorescence Imaging

Immunofluorescence images were acquired using a QICAM Fast 1394 Cooled Digital Camera, 12-bit, Monochrome CCD camera (4.65 x 4.65 µm pixel size and 1.4 MP, 1392 x 1040 pixels) using Micromanager and a 40x objective lens on a Nikon TE2000 inverted widefield fluorescence microscope. Cells were imaged using transmitted light to find the optimal focus on the field of view to observe the nuclear blebs. Ultraviolet light (excitation 360 nm) was used to visualize DNA via DAPI, blue fluorescent light (excitation 480 nm) was used to visualize lamin B1, and green fluorescent light (excitation 560 nm) was used to visualize emerin. Green fluorescent light (excitation 560 nm) was also used to visualize γH2AX. Images were saved and transferred to NIS-elements (Nikon) or FIJI (Schindelin et al., 2012) for analysis.

Immunofluorescence images of cells that were previously time lapsed were acquired using a Nikon Instruments Ti2-E microscope with Crest V3 Spinning Disk Confocal, Hamamatsu Orca Fusion Gen III camera, Lumencor Aura III light engine, TMC CleanBench air table, with 40x air objective (N.A 0.75, W.D. 0.66, MRH00401), and a 12-bit camera through Nikon Elements software. Images were taken at 0.5 µm z-steps over 4.5 µm. Ultraviolet light (excitation 408 nm) was used to visualize DNA via DAPI, green fluorescent light (excitation 546 nm) was used to visualize emerin or lamin B1, and red fluorescent light (excitation 638 nm) was used to visualize γH2AX or laminB1.

### Nuclear Bleb Analysis

As outlined previously (Bunner et al., 2024), images were analyzed in NIS-elements or exported to FIJI (Schindelin et al., 2012) to analyze the intensity of each component by normalizing bleb intensity to nuclear body intensity. The body, bleb, and background of each nucleus image were measured by drawing regions of interest (ROI) via the polygon selection tool. Measurements of mean intensity for each ROI were recorded and exported to Microsoft Excel. Within Excel, the background intensity was subtracted from the body and bleb. Then the average bleb intensity was divided by the average nuclear body intensity to give a relative measure, where the same average intensity would result in 1. These values were transferred to Prism where Mann-Whitney test, unpaired t-tests, or one-way ANOVA were performed, and the data was graphed: not sure this is accurate.

### DNA Damage Analysis

To analyze γH2AX data for presence of DNA damage, the image background intensity was subtracted and the Cy5 channel was selected to observe γH2AX foci in the nuclei. On Nikon NIS-elements or FIJI, binary bright spot detection was used to record γH2AX foci. A diameter of 0.5 µM was used to count foci. Data was exported to an Excel file where an average number of foci for each treatment condition was calculated.

### Live cell time lapse fluorescence imaging and analysis

As previously described, we used either established MEF NLS-GFP stable cell lines to quantify nuclear shape and rupture (Berg *et al*., 2023; Pho *et al*., 2023). Images were acquired with Nikon Elements software on a Nikon Instruments Ti2-E microscope, Orca Fusion Gen III camera, Lumencor Aura III light engine, TMC CleanBench air table, with 40x air objective (N.A 0.75, W.D. 0.66, MRH00401). Live cell time lapse imaging was possible using the Nikon Perfect Focus System and Okolab heat at 37°C, supplemental humidity, and a 5% CO2 stage top incubator (H301). Cells were imaged in single well gridded cover glass dishes (Cellvis #D35-14-1.5GI). For time lapse data, images were taken in 2-minute intervals for 3 hours over 10 fields of view. Images were captured at 50ms exposure time, 12-bit depth, and 4% blue fluorescent light (475 nm) power. Time lapse images were analyzed by visually tracking nuclear bleb formation and rupture throughout the movie. The frequency and recency of nuclear ruptures and bleb formation and lifetime were recorded. To determine bleb lifetime, the time of bleb formation was subtracted from the total duration of the time lapse.

LNCaP nuclei were transiently transfected with NLS-GFP via the CellLight Nucleus-GFP (ThermoFisher Scientific, C10602). Cells were transfected with 50 or 100 µL of CellLight two days prior to imaging.

## Acknowledgements

We would like to thank HHMI which purchased microscopes used in Bioimaging class via a grant and The Biology Department at UMass Amherst for the use of the ISB 360 facilities.

## Funding

This work was primarily supported by NIH NIGMS grant Maximizing Investigators’ Research Award R35GM154928.

### Data availability

All raw data that is available https://doi.org/10.6084/m9.figshare.28451933. Image data set can be made available upon request.

### Competing interests

The authors declare no competing interests.

## Notes

### Competing Interest Statement

The authors have declared no competing interest.

