## Supplemental Figure 1 for "Lamin B loss in nuclear blebs is rupture dependent while increased DNA damage is rupture independent"

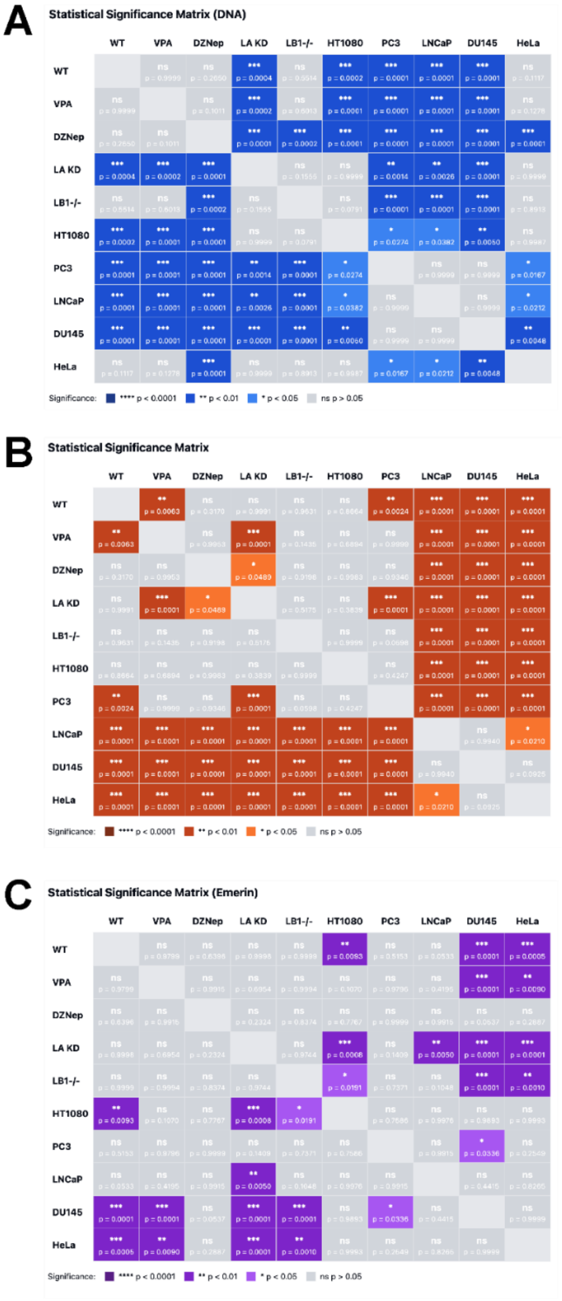


**Supplemental Figure 1. Statistical Analysis of Nuclear Bleb Composition Across Cell Types and Perturbations.** Complete statistical significance matrices from one-way ANOVA with Tukey's post-hoc test for nuclear bleb/body ratios of DNA (A), lamin B (B), and emerin (C) across all experimental conditions. Analysis includes MEF wild type (WT), chromatin perturbations (VPA, DZNep), lamin perturbations (LA KD, LB1-/-), and human cancer cell lines (HT1080, PC3, LNCaP, DU145, HeLa). Each panel displays results as a heat map where color intensity corresponds to significance level: DNA (blue shades), lamin B (orange shades), and emerin (purple shades), with darker shading indicating higher significance (***p < 0.001 , **p < 0.01, *p < 0.05), while grey indicates non-significant comparisons (ns, p > 0.05). Each matrix element shows the exact p-value for the corresponding pairwise comparison between conditions."
